# American mink as an animal model to study SARS-CoV-2 and vaccine response

**DOI:** 10.1101/2025.06.10.658224

**Authors:** Kirsi Aaltonen, Jenni Virtanen, Kristel Kegler, Vinaya Venkat, Lauri Kareinen, Essi M. Korhonen, Thanakorn Niamsap, Rasmus Malmgren, Johanna Korpela, Rauno A. Naves, Maarit Iso-Oja, Sofie Svenns, Tomas Häggvik, Eeva Ojala, Olli Vapalahti, Jussi Peura, Antti Sukura, Olli Ritvos, Arja Pasternack, Heli Nordgren, Ravi Kant, Tarja Sironen

**Affiliations:** Faculty of Veterinary Medicine, Department of Veterinary Biosciences, University of Helsinki, Helsinki, Finland; Faculty of Medicine, Department of Virology, University of Helsinki, Helsinki, Finland; AnaPath Services GmbH, Switzerland; Finnish Food Authority, Helsinki, Finland; Faculty of Biological and Environmental Sciences, Molecular and Integrative Biosciences Research Programme (MIBS), University of Helsinki, Helsinki, Finland; Fevia Fin Lab Oy Ab, Vaasa, Finland; Nordic SARS Response Ab; Kannus Research Farm Luova Ltd., Kannus, Finland; Kronovet Ab, Kruunupyy, Finland; Helsinki and Uusimaa hospital district, HUSLab, Helsinki, Finland; Finnish Fur Breeders’ Association FIFUR, Finland; Finnish Centre for Laboratory Animal Pathology (FCLAP), Helsinki Institute of Life Science (HiLIFE), University of Helsinki, Helsinki, Finland; Faculty of Medicine, Department of Physiology, University of Helsinki, Helsinki, Finland; Department of Tropical Parasitology, Institute of Maritime and Tropical Medicine, Medical University of Gdansk, 81-519 Gdynia, Poland

**Author notes:** These authors contributed equally to this work (Kirsi Aaltonen, Jenni Virtanen, Kristel Kegler). Corresponding author (KA).

## Abstract

Selecting a suitable animal model is crucial in understanding infectious diseases and developing vaccines. Here, we developed a receptor-binding domain -based SARS-CoV-2 vaccine with mouse Fc an immunopotentiator in a mink model. Four different variations of the vaccine were tested in groups of 30-31 American mink and followed for IgG and neutralizing antibodies (nAb) up to 27 weeks. Subcutaneous version induced a strong IgG and nAb response within two weeks and was still detectable at 27 weeks. Intranasal version also caused a detectable, although weaker, immune response. A simultaneously given subcutaneous vaccine against virus enteritis, botulism and hemorrhagic pneumonia potentially caused a lower SARS-CoV-2 antibody response, highlighting the need for further studies on co-effects of vaccines. In virus challenge with Alpha variant (B.1.1.7), vaccinated mink had a stronger antibody response than unvaccinated mink. Despite not preventing the infection, vaccinated mink had milder clinical signs and less virus in saliva. Another challenge of unvaccinated mink with Omicron variant showed similar results to alpha (BA.1) variant. Virus RNA was detected in the brain of unvaccinated mink but not vaccinated mink by *in situ* hybridization, indicating a suitability of mink to study neurological effects of SARS-CoV-2 and potentially long COVID as well.

**Author summary:** Finding a good animal model is very important in studying infectious diseases and their treatment. The recent SARS-CoV2 pandemic highlighted this dilemma. We have developed an animal model based on mink due to their close match to humans in the symptomology of this disease. Such model will enable the study of relative susceptibility, transmission, tissue tropism, complex pathogenesis and long COVID, as well as prophylaxis and vaccines. This model will also help reduce the use of primates in this research.

We have further developed a new vaccine for SARS-CoV2 based on the receptor binding domain of the S-protein with an inbuilt immune-enhancer. This vaccine underwent extensive testing in mink to determine response, long term protection, and safety. The vaccine was found to provide excellent titers of neutralizing antibodies with wide range in target variants. It also reduced the severity and duration of visible symptoms in the animals significantly. We propose this vaccine candidate for further study and future commercialization.

## Introduction

Severe acute respiratory syndrome coronavirus 2 (SARS-CoV-2) is the third highly human pathogenic member of family *Coronaviridae* to emerge in the 21^st^ century(1). Due to constant virus evolution, new threatening variants can still emerge, justifying WHO’s efforts in monitoring virus mutations in animals and humans to aid in vaccine development(2, 3). Additionally, long COVID has been estimated to affect tens of millions of people around the world(4). To understand infectious diseases in both human and animal hosts and to develop vaccines, special attention should be paid to selecting an appropriate animal model. In SARS-CoV-2 research and vaccine development, hamsters, mice, ferrets, and non-human primates have been used; each model has advantages and limitations(5). When it comes to long COVID, golden hamsters have been suggested(6). However, more information is needed for selecting the best animal model for each purpose, especially when it comes to virus entrance into the brain and the long-term effects of COVID-19.

Mink is a favorable animal model for COVID-19 due to the clinical picture similar to humans(7, 8). The virus replicates efficiently in the upper and lower respiratory tracts and can cause acute, severe interstitial pneumonia or diffuse alveolar damage with varying clinical signs, such as labored breathing, nasal discharge, and anorexia(8, 9). The virus is most readily detected in throat and nasal samples but is also secreted to feces in both symptomatic and asymptomatic mink, indicating also asymptomatic spread of the virus(9, 10). A spike protein-based subunit vaccine may prevent replication in the upper and lower respiratory tracts and limit SARS-CoV-2-induced lung damage, but studies with different variants and larger sample sizes are required(8).

SARS-CoV-2 is known to jump species-humans to other animals and vice versa-raising concerns about new zoonotic mutants spreading among humans(11). Spread of the virus among farmed mink and observations in wild mink adds to this growing concern(10, 12–18). In addition to causing epidemics among mink, spillover of mink-related variants back to humans has been reported(15), causing the Danish National Institute of Public Health announced the culling of all 17 million mink in the country. The data available from Denmark on these mink-associated SARS-CoV-2 variants suggest that these variants can spread rapidly in both the mink farms and nearby human communities, increasing the need to understand SARS-CoV-2 in mink(19). Hence ECDC (European Centre for Disease Prevention and Control) and WHO have urged the surveillance of the host-animal interface for early detection for VOCs (Variants Of Concern)(2, 20).

In this study, we developed a vaccine against SARS-CoV-2 using mink as an animal model. We also performed a virus challenge in both vaccinated and unvaccinated animals to demonstrate its effectiveness in immune response induction and symptom control.

## Results

### Vaccination against SARS-CoV-2 induces high IgG levels and NAb titers in mink

One objective was to develop a vaccine against SARS-CoV-2 based on the receptor binding domain (RBD) of the Wuhan strain (MN908947.3) with American mink as an animal model. Mouse Fc (mFc) was included to determine if this would enhance the immune response. Four versions were tested in groups of 30-31 animals. Two received either 40 µg (RBD mFc big) or 10 µg (RBD mFc small) of RBD with mFc subcutaneously, one group (RBD small) received 10 µg of subcutaneous RBD without mFc and one intranasal RBD with mFc (40 µg). A control group received alum adjuvant (Alhydrogel) only. A second dose was given to all groups 2 weeks after the initial inoculation.

Vaccine responses were monitored by collecting blood every 2-4 weeks and testing IgG levels with enzyme-linked immunosorbent assays (ELISA, Fig. 1a-b, Table S1). An increase in antibody levels was detected as early as 2 weeks after the first dosing in all mink in RBD mFc big, RBD mFc small, and RBD small groups. The levels rose either close to or above the upper detection limit by week 4. The highest levels were observed in mink vaccinated with the larger dose of RBD mFc. Antibody levels remained high through the follow-up period (11 weeks from the first dose for RBD mFc small and RBD small or 27 weeks for RBD mFc big and intranasal), although levels started to slowly decline 5-10 weeks after the first dose. Elevated IgG levels were also detected in 32% (9/28) mink that received an intranasal vaccination. Five mink showed elevated ELISA values (0.422-0.931) in the pre-vaccination sample and were hence tested with a microneutralization assay (MN). No neutralizing antibodies (nAbs) were detected. Before the experiment, a panel of mink samples from before the pandemic were tested to validate the protocol and the false positive rate was 1% (supplementary data).

**Fig. 1.**
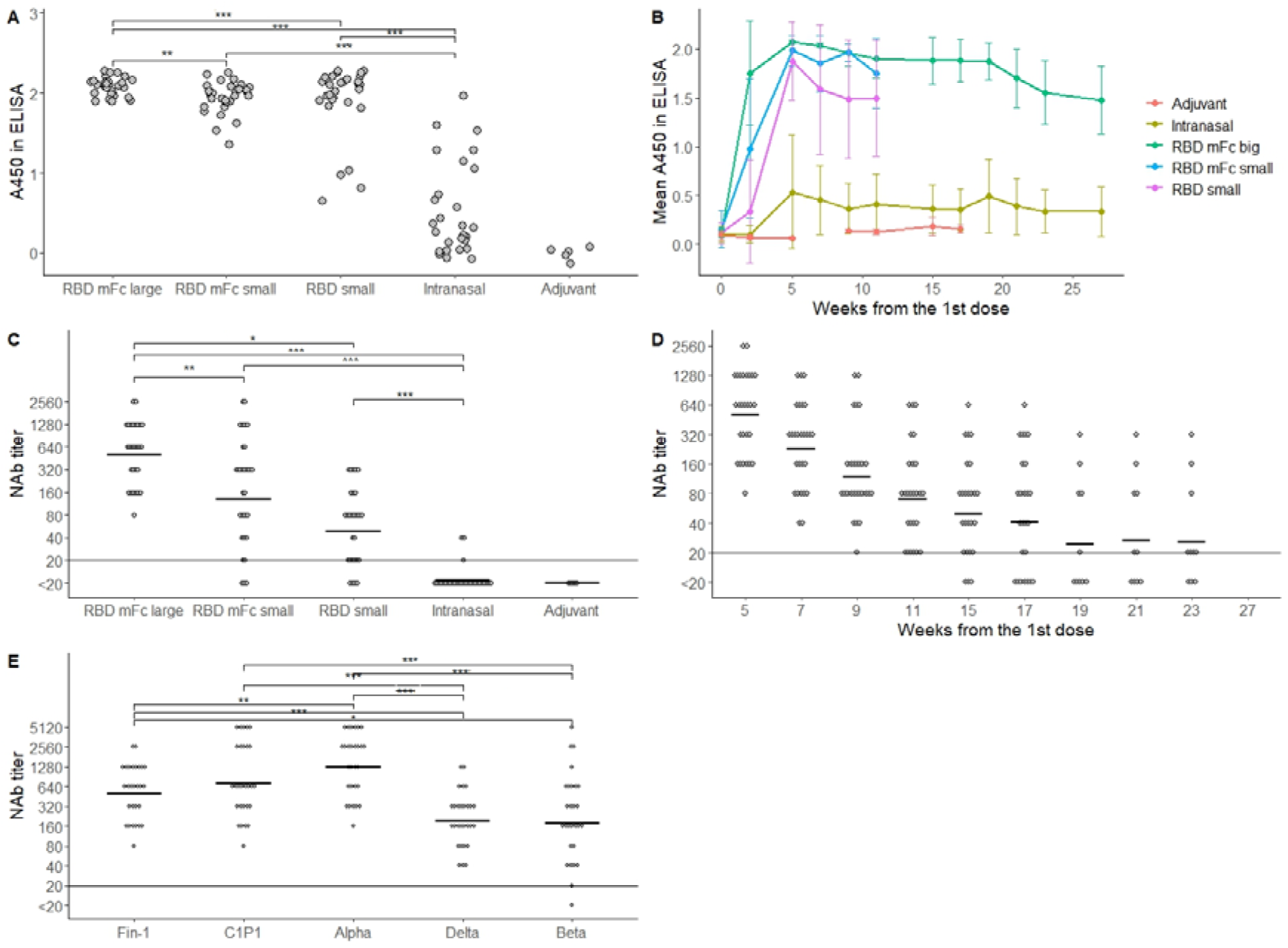
Antibody responses against SARS-CoV-2 after vaccination. (a) IgG levels (A450 in ELISA) of different groups 5 weeks after first dose and 3 weeks after booster. (b) Follow-up of IgG levels after vaccination (error bars as +/- SD). (c) nAb titres of different vaccine groups against FIN-1 strain 5 weeks after first dose. (d) Follow up of nAb titres of mink vaccinated with RBD mFc big. (e) nAb titres against different variants in RBD mFC big group 5 weeks after first dose. The upper limit of detection is 2.0-2.1 in ELISA. nAb titres are expressed in log_2_ scale and limit of detection and GMT-values are marked with horizontal lines. Statistically significant differences (*: <0.05, **: <0.01, and ***: <0.001) are indicated in a, c, and e.

Blood samples from all groups were tested for nAbs with MN 5 weeks after the first dose (Fig. 1c). Geometric mean titers (GMT) were 548 for RBD mFc big, 175 for RBD mFc small, 58 for RBD small, and 11 for intranasal vaccination. Titers were below detection limit (<20) in all control mink and most intranasally vaccinated mink, except for 3 animals with low titers between 20-40. Although GMTs were slightly lower in males than females (approximately 1.5-2.0-fold difference) in all groups, the differences were not statistically significant (p-values 0.069-0.486) (Fig. S1 and Table S2). Additionally, the RBD mFc big group was analyzed for nAbs at nine timepoints to follow the decline in nAb levels (Table S3). GMT was highest (538) 5 weeks after the first dose and decreased to 30 by week 23 (Fig. 1d). When different virus variants were compared (Fig. 1e), high titers were detected with the wild-type strain C1P1 and the Alpha variant, whereas titers against Beta or Delta were smaller but still detectable 5 weeks after the first dose.

### Other vaccinations given simultaneously may hinder the immune response to SARS-CoV-2

We examined if Febrivac 3-Plus vaccine (inactivated mink enteritis virus, type C botulinum toxoid, and inactivated *Pseudomonas aeruginosa*) - routinely given to mink at the age of 6-8 weeks - would affect SARS-CoV-2 vaccine response when given simultaneously. For the experiment, 20 mink were vaccinated with 40+40 µg of RBD mFc and 20 with 10+10 µg of RBD mFc. From both groups, 10 animals were vaccinated with Febrivac 3-Plus and the SARS-CoV-2 vaccination simultaneously, and 10 animals were inoculated 5 weeks after the first dose of the SARS-CoV-2 vaccine (Table S4).

IgG levels and nAbs were analyzed from blood samples taken 5 weeks after the first dose (Fig. S2). Due to several values being above the detection limit of 2.0-2.1, samples from mink that received the large dose were analyzed at 1:1000 dilution in ELISA. In these mink, mean absorbance (A450) in ELISA was 1.877 (SD = 0.165) when the SARS-CoV-2 vaccine was given at the same time with Febrivac 3-Plus and 1.961 (SD = 0.069, p=0.481, Table S5) when there was a gap of a couple of weeks between the vaccines. In MN, GMTs were 235 (simultaneous vaccination) and 549 (vaccines administered separately, p = 0.190). In the RBD mFc small group, the corresponding GMT values were 1.879 (SD = 0.113) and 2.010 (SD = 0.018, p = 0.001) in ELISA and 98 and 260 (p = 0.063) in MN, respectively. The group with the greatest IgG and nAb response (RBD mFc big, Febrivac 3-Plus given 5 weeks after first dose) was analyzed for nAb against the Omicron variant. A lower, but measurable response (GMT = 34) was observed.

### Antibody response, virus levels and clinical signs under virus challenge differ between vaccinated and unvaccinated animals

The group vaccinated with RBD mFc big was selected for a virus challenge with the Alpha variant. Fifteen mink vaccinated with the first dose 5-6 months earlier were selected. Additionally, 5 unvaccinated mink that had received adjuvant only were selected as controls. Infected mink were followed for 7 days, after which they were sacrificed and examined for pathology.

Mean IgG levels in unvaccinated mink increased from a mean of 0.13 (SD 0.03, considered negative) to a mean of 1.74 (SD 0.19) (Fig. 2a, Table S6). IgG levels before infection were above the upper limit of detection in 5/15 vaccinated mink; the mean value for the other mink was 1.55 (SD 0.4). After challenge, IgG levels were above 2.0 in all vaccinated mink. None of the unvaccinated mink had nAbs before infection and GMT after challenge (Fig. 2b) was 243. GMT of vaccinated mink was 139 before infection. After infection, all vaccinated mink had titers >2560.

**Fig. 2.**
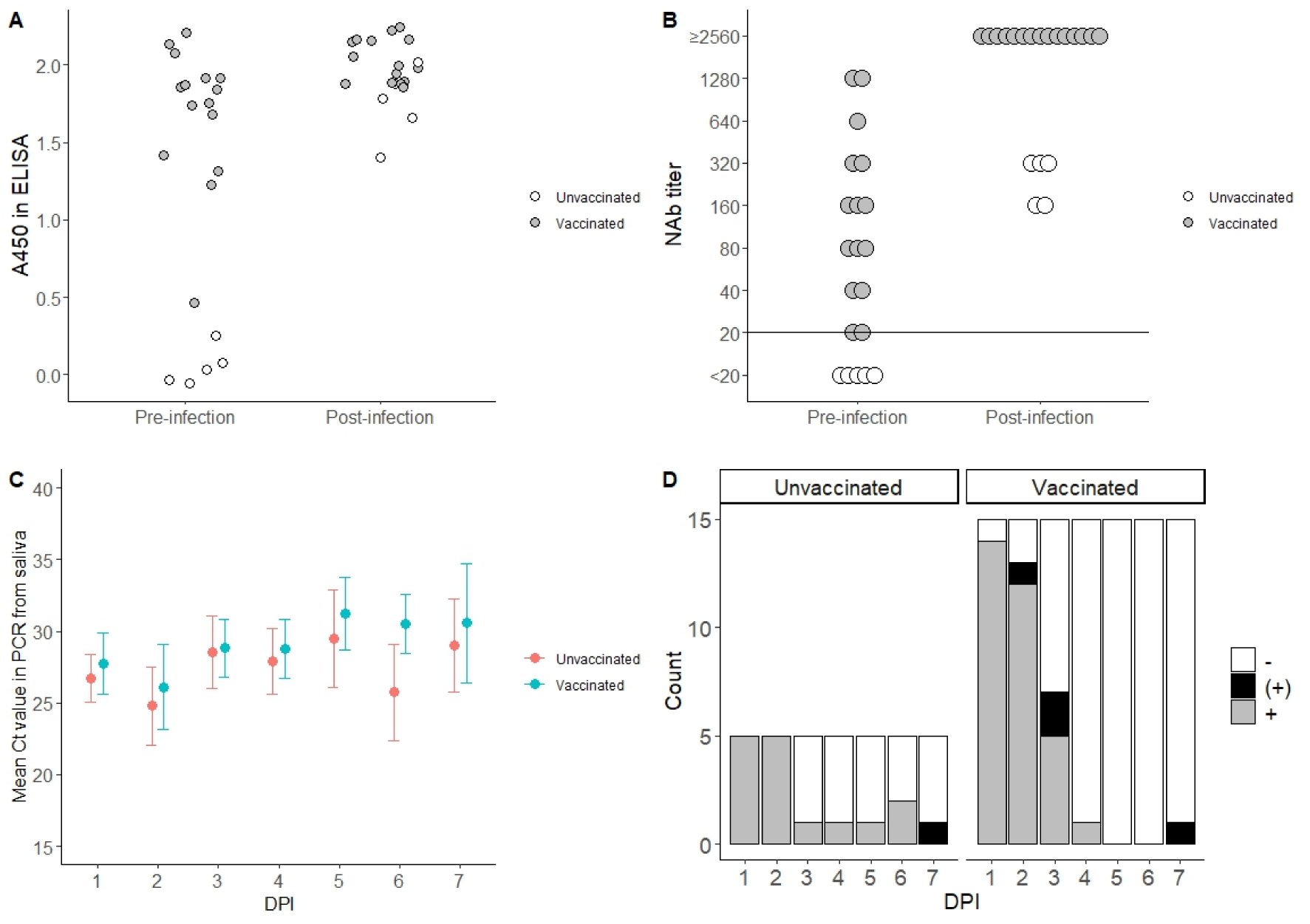
Antibody response, PCR, and culture results in virus challenge with Alpha variant in vaccinated and unvaccinated mink. IgG levels (A450 in ELISA) (a) and nAb titre (b) pre- and post-infection, Ct value in RT-qPCR from saliva (c), and cell culture results (d). ELISA upper detection limit is 2.0-2.1 and MN lower limit of detection is marked with horizontal line. Culturing results: - = negative, (+) = possible positive, and + = positive.

Virus was consistently detected in saliva by PCR. Ct values slowly increased by the end and were higher in vaccinated mink (p = 0.007) (Fig. 2c). No difference was detected between genders (p = 0.801). Infectious virus was detected in the saliva of all unvaccinated mink and in most of the vaccinated mink for the first 2 days, after which the proportion started to decrease. Although virus was cultured from vaccinated mink until 6 days post-infection (dpi) and from unvaccinated mink until 4 dpi, this was not statistically significant (p = 0.389) (Fig. 2d).

Infected mink exhibited varying degrees of nasal irritation (runny or dry nose and sneezing), diarrhea, anorexia, and malaise (Table S7). Runny nose and sneezing were observed more in unvaccinated mink (mean score = 5.00) than vaccinated mink (mean = 2.13). Differences were smaller with diarrhea (unvaccinated = 3.60 and vaccinated = 2.40), anorexia (unvaccinated = 3.40 and vaccinated = 2.73), eye symptoms (unvaccinated = 0.20 and vaccinated = 0.13), and malaise (unvaccinated = 0.60 and vaccinated = 0.20). Mild cough was observed only in 1 vaccinated mink on 2 days (mean = 0.13). Clinical signs were detected slightly more in males than females in both groups (Fig. S3).

### Histopathological lesions in the upper and lower respiratory tracts were milder in vaccinated mink after infection with Alpha variant

All mink had neutrophil-rich mucinous secretions on the nasal conchae. The nasal mucosa exhibited loss of cilia, multifocal swelling, and degeneration and desquamation of epithelial cells (Fig. 3a, Tables 1 and S8). The severity of lesions varied in the respiratory and olfactory region from mild to severe. Immunohistochemistry revealed free SARS-CoV-2 antigen in the nasal mucinous secretions in all mink. Multifocal or focal virus antigen was observed in the epithelial cells of the respiratory and olfactory region in all unvaccinated mink. One mink expressed viral antigen also in the submucosal glands in the respiratory region and two mink in the gland and lymphoid tissue of the olfactory region. In contrast, only 3/15 vaccinated mink exhibited viral antigen focally in the respiratory region and 7/15 in the olfactory region epithelial cells. Viral antigen was detected in lymphoid tissue in three vaccinated mink (3/15).

**Fig. 3.**
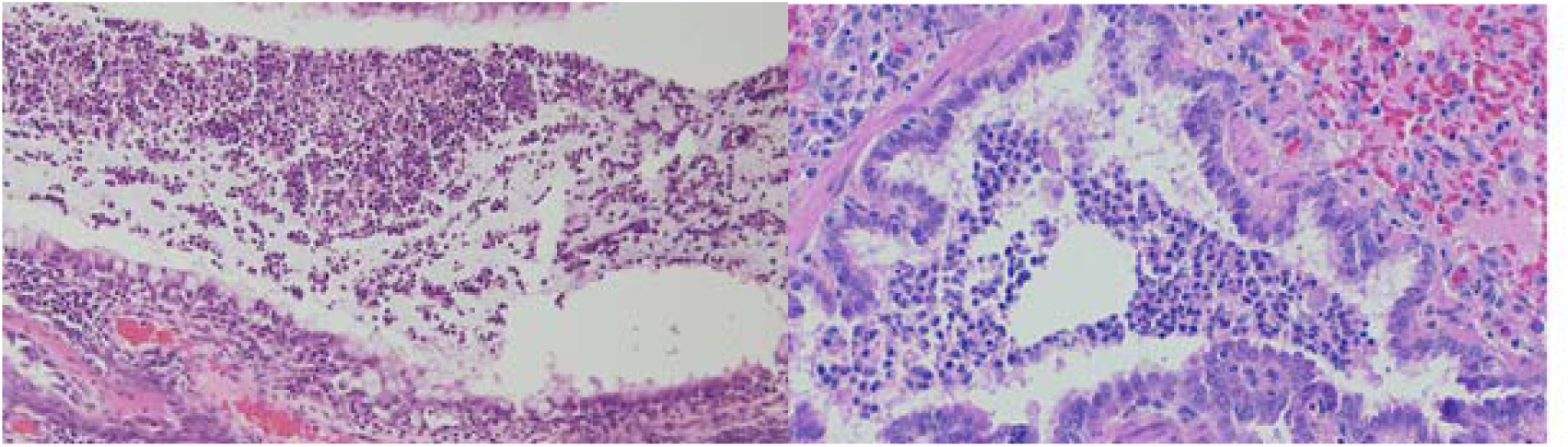
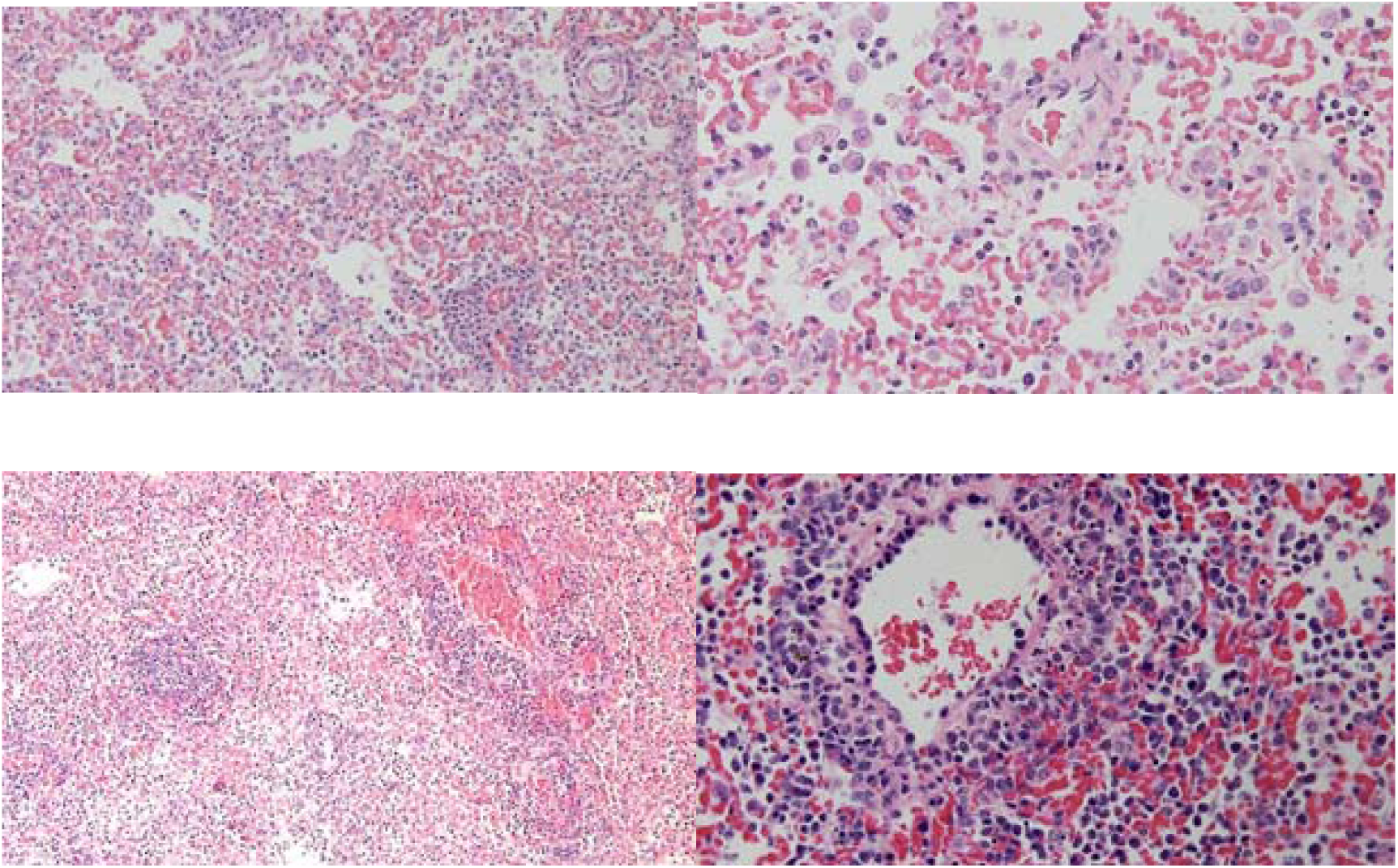
Histopathology of nasal cavity and lung of unvaccinated mink experimentally infected with SARS-CoV-2. (a) Nasal cavity, respiratory region. Suppurative secretion in the nasal cavity, loss of cilia, mucinous and multifocal swelling, and degeneration and desquamation of epithelial cells. Haematoxylin and eosin (HE). (b) Lung. Neutrophil-rich secretions in the bronchiole, some neutrophils have infiltrated to the bronchial epithelium, syncytial formation (HE). (c) Lung. Diffuse interstitial pneumonia, alveolar damage, and increased cellularity in the alveoli (HE). (d) Alveolar damage and pneumocytes and alveolar macrophages in the alveoli. (e) Lung. Oedema and mononuclear cells around blood vessels (HE). (f) Lung. Perivascular mononuclear inflammatory cells and infiltration to the vessel wall (fibrinoid arteritis) (HE).

Unvaccinated mink had histopathological lesions in the lungs that resembled those described in natural and experimental SARS-CoV-2 infections in mink(8, 9). Trachea, bronchi, and bronchioles exhibited neutrophil-rich mucinous exudate, mild to moderate multifocal flattening of epithelial cells, loss of cilia, multifocal degeneration of the epithelium, and formation of syncytial giant cells (Fig. 3b). The main histopathological finding in parenchyma was multifocal to coalescing bronchointerstitial pneumonia. The alveolar septae were thickened by oedema and by lymphocytes and to a lesser extent by neutrophil infiltrates.

Activation and hyperplasia of type II pneumocytes, foamy intra-alveolar macrophages, lymphocytes and neutrophils, and small fibrin deposits indicated acute alveolar damage (Fig. 3c, d). Varying degrees of perivascular oedema and perivascular inflammatory cell infiltration, endothelial activation and, occasionally, true vasculitis were present (Fig. 3f). Vaccinated mink showed qualitatively similar but quantitatively milder and fewer lesions in the lungs (Fig. 4). In particular, formation of syncytial cells, alveolar damage, and vascular lesions were not as evident as in unvaccinated mink (Table 1). Lesion scores of unvaccinated infected mink were severe in 60% (3/5) mink and moderate in 20% (2/5) mink; the corresponding values in vaccinated animals were 7% (severe, 1/15) and 27% (moderate, 4/15). All (5/5) unvaccinated mink exhibited SARS-CoV-2 antigen in the lungs in immunohistochemistry, however, the number of positive cells was very low in 3 mink. All unvaccinated mink exhibited viral antigen in focal or multifocal groups of bronchial or bronchiolar epithelial cells, two mink exhibited viral antigen also in pneumocytes. In contrast, viral antigen was detected only in a few cells or in a single cell in the bronchial epithelium in 5/15 or pneumocytes in 2/5 vaccinated mink. No viral antigen was detected in the uninfected vaccinated or unvaccinated control animals.

**Fig. 4.**
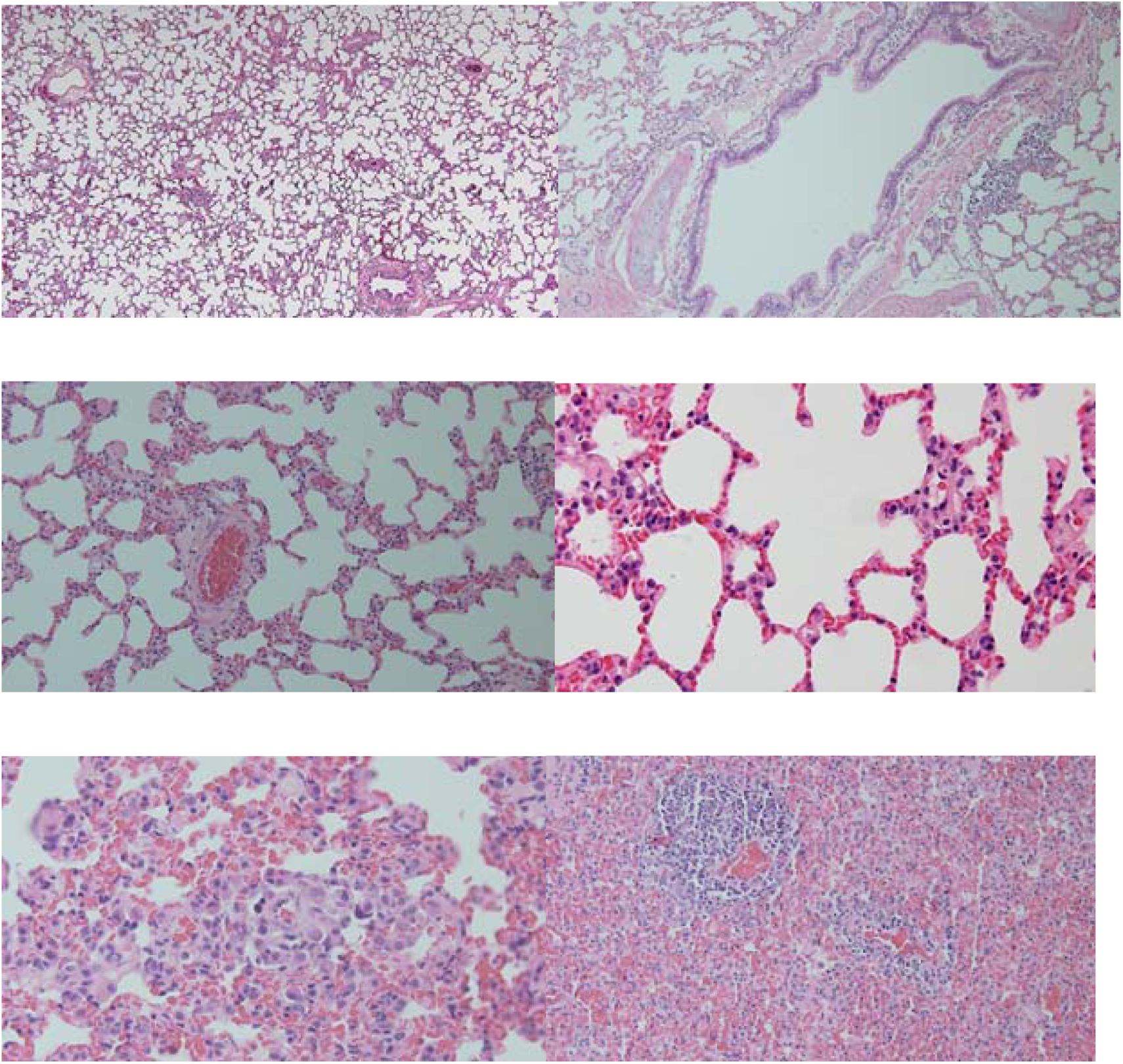
Lung histopathology of vaccinated mink experimentally infected with SARS-CoV-2. (a, b, and c) Lung. Lung parenchyma, bronchi, and blood vessel with no specific histopathological lesions. Haematoxylin and eosin (HE). (d) Lung. Very mild pneumocyte II activation (HE). (e) Moderately increased number of inflammatory cells in the interstitial lung tissue, moderate pneumocyte II activation (HE). (f) Moderate vascular lesion, congestion, and alveolar oedema (HE).

**Table 1.**
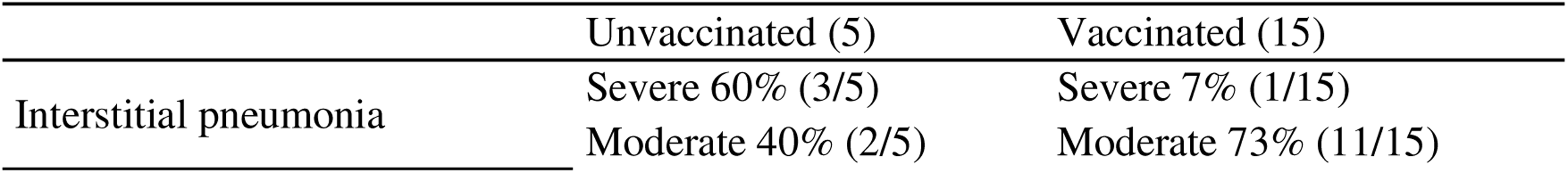
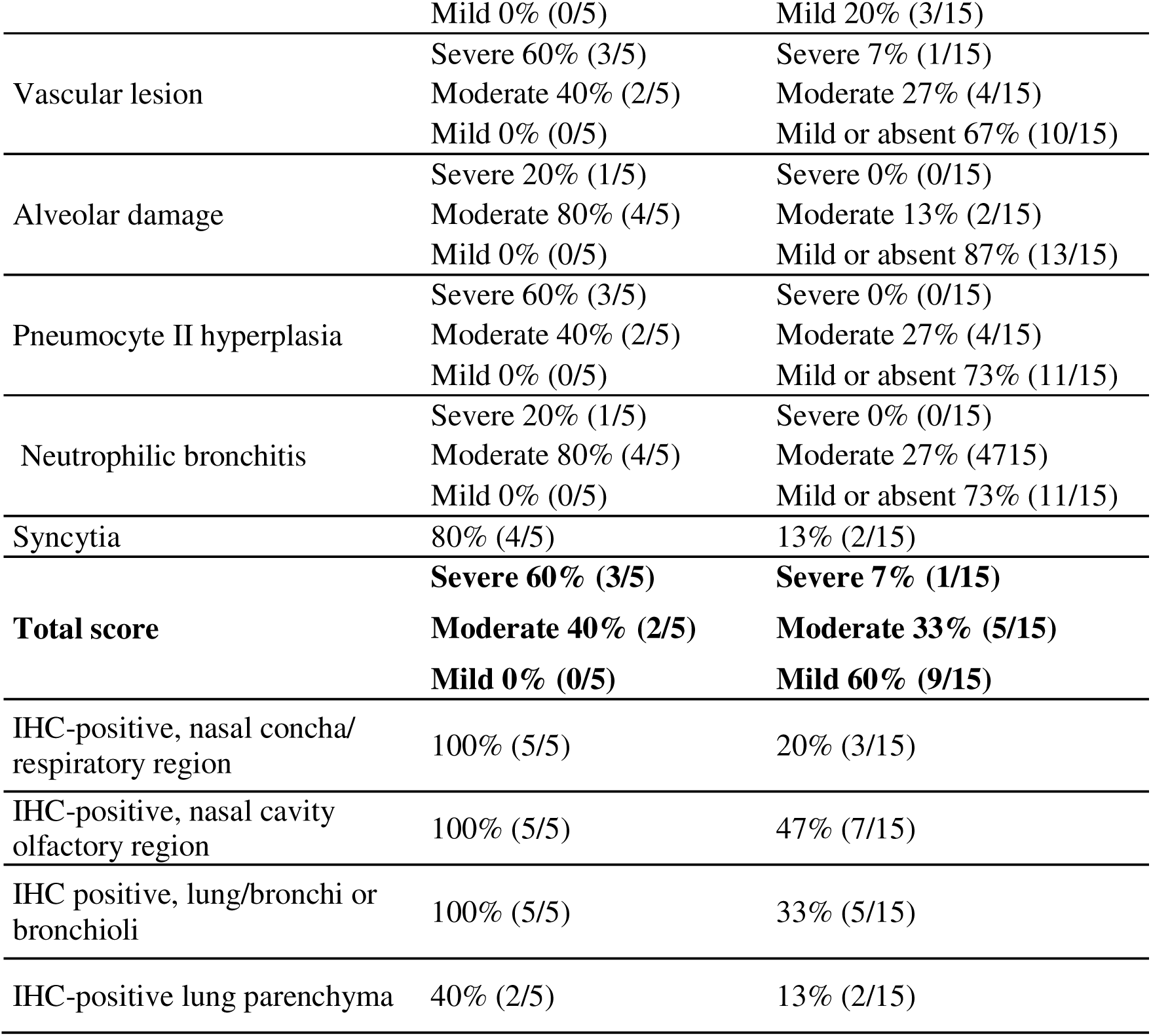
Histopathology in nasal cavity and lungs of unvaccinated and vaccinated SARS-CoV-2 infected mink. The total score of lung lesions is based on severity of bronchitis/bronchiolitis, interstitial pneumonia, alveolar damage, vascular lesions, and degree of pneumocyte type II hyperplasia.

### Vaccination against SARS-CoV-2 prevents virus dissemination to the central nervous system

We explored SARS-CoV-2 distribution in the central nervous system (CNS) of unvaccinated and vaccinated mink after viral challenge with the Alpha variant. Brains were transected at the levels of the olfactory bulb, frontal lobe, optic chiasm, pretectal region (thalamus), trochlear nucleus, pons, and cerebellum and subject to histopathologic evaluation and *in situ* hybridization (ISH) for SARS-CoV-2 RNA detection. Additionally, bilateral sections of the trigeminal ganglia and trigeminal nerves were included.

Despite the lack of histomorphologic changes in the brain, SARS-CoV-2 RNA was detected in all unvaccinated mink (n=5). Four animals had low, and one animal had moderate viral RNA load. Positive hybridization was strictly observed in the cortical grey matter within the cytoplasm (soma) of cells morphologically compatible with neurons. No hybridization was present in the white matter, nor in endothelial cells or other structures from vascular walls, ependymal cells, and meninges.

Viral RNA was distributed in the frontal cortex, premotor cortex, primary motor cortex, primary and secondary somatosensory cortex, caudate nucleus, cingulate cortex, posterior parietal cortex, suprasylvian cortex, lateral entorhinal cortex, hippocampus, and caudal colliculus (Fig. 5). SARS-CoV-2 RNA hybridization was not observed in the olfactory bulb, pons, cerebellum, trigeminal ganglia, and trigeminal nerves. Surprisingly, all vaccinated mink (n=15) lacked viral RNA in the CNS.

**Fig. 5.**
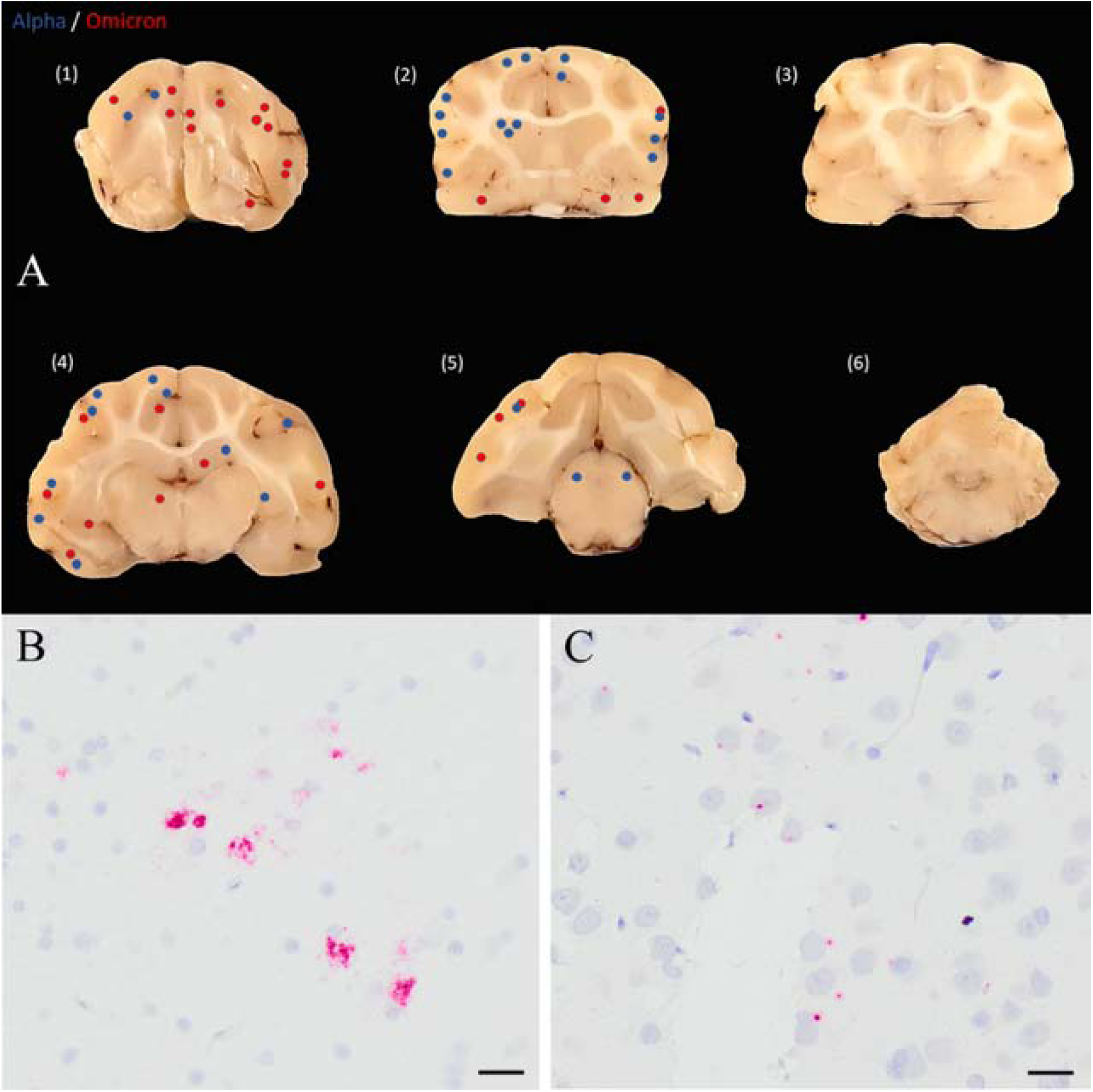
Comparative CNS infection in mink infected with Alpha and Omicron variants. (a) Distribution of SARS-CoV-2 within the brain of unvaccinated mink infected with Alpha and Omicron variants. Transversal sections of the brain at the levels of (1) frontal lobe, (2) optic chiasm including basal nuclei, (3) caudal diencephalon, (4) pretectal region (thalamus), (5) midbrain (pons), and (6) cerebellum and brain stem. The dots represent the presence of the virus within the brain without distinction of each individual animal. (b) SARS-CoV-2 RNA S-gene probe hybridization (red) is prominent within the neuronal bodies in the cortex of transection level 2 (optic chiasm and basal nuclei) of an unvaccinated mink experimentally infected with the Alpha variant. Bar 50 µm. (c) Probe hybridization in transection level 1 (frontal cortex) of a mink experimentally infected with Omicron variant. Note the presence of SARS-CoV-2 RNA (red) within neuronal bodies and within neuronal nuclei. Bar 20 µm.

### Cross comparison of Alpha and Omicron strains between male and female mink

Previously, we experimentally infected three male American mink with the Omicron variant and left two mink as uninfected recipients and observed symptomatic disease and mink-to-mink transmission in these animals(21). Here, we repeated the experiment with four female mink to compare the two genders and two variants.

Similar to previous experiment with male mink(21), Omicron-infected saliva from female mink were PCR positive from 1 dpi and remained positive over 7 days (Table S9 and S10). One recipient mink was PCR positive every day from 1 dpi and another was PCR positive at 1, 8, and 9 dpi. Both infected mink were cell-culture positive 1 dpi but subsequently remained generally negative. One of the recipients was culture positive at 7 dpi (1 day before turning PCR positive); recipients were otherwise generally culture negative.

Infected mink exhibited a variety of clinical signs, including malaise (lethargy, crouching, fluffiness, heavy breathing), signs of nasal irritation, cough, anorexia, diarrhea, and watery or crusty eyes (Table S11). In the acclimation period before infection, poor appetite, mild diarrhea, and occasional watery eyes were observed. No clear difference between genders was observed; some signs were more common in males and some in females (Fig. S3b). Although most clinical signs were more common in recipients than in experimentally infected mink, recipients were followed for 3 days longer, and thus direct comparison is difficult (Fig. S3c).

Overall, microscopic lesions in omicron-infected mink were comparable to animals inoculated with the Alpha variant, as described above (Table S12). Both omicron-inoculated female mink had identical, but slightly milder histologic changes in the upper and lower respiratory tracts as did male mink(21). In these two animals, viral nucleoprotein was variably distributed in the lumen and mucosal cells from the respiratory epithelium, focally present in the olfactory mucosa, and observed within the affected pulmonary parenchyma in 1/2 mink. Recipient female mink had very mild microscopic lesions in the respiratory and olfactory mucosa, and mild to moderate changes in the lungs. Focal viral antigen expression was detected in the respiratory mucosa in both animals, and focally in the olfactory region in 1/2 mink. Viral antigen was focally present in the lung parenchyma in both recipient mink. In contrast to animals infected with Alpha variant, bronchi and bronchiole in Omicron-infected female and male mink lacked viral antigen expression.

The distribution of SARS-CoV-2 RNA in the CNS of female and male mink infected with Omicron variant was identical to the one observed in animals infected with Alpha variant, with the exception than in one inoculated mink, SARS-CoV-2 RNA was present in axons of the trigeminal ganglia. Interestingly, male and female recipient mink also had detectable viral RNA in the brain. In contrast to animals infected with Alpha variant, omicron-infected mink had SARS-CoV-2 RNA not only in the neuronal cytoplasm but also within the nucleus and near the nuclear membrane.

## Discussion

We developed an RBD-based SARS-CoV-2 vaccine and tested it in mink. Vaccine was shown to induce a strong immune response that decreased the clinical sings and viral load in saliva months after vaccination and prevented virus entrance into the brain. Additionally, we showed that the infection in experimentally infected mink mimics human infection, also when it comes to neurotropism, indicating that mink is a good animal model to understand not only SARS-CoV-2 but possibly long COVID as well.

As most nAbs against SARS-CoV-2 target RBD, the RBD has been suggested as a potential vaccine target(22–24). Fc fragments may act as an immunopotentiator by promoting interaction of the vaccine with Fc receptors on antigen-presenting cells and thus prolong the nAb response. This approach has shown good results in transgenic mice(25). Our results with mink support earlier studies conducted in other lab animals. Although an effective IgG and nAb response was observed in mink vaccinated with RBD with and without mFc, a significantly stronger response was observed when mFc was included in the vaccine. Based on these results, Fc fragments may have considerable potential in vaccine development. The 40-µg dose provided a significantly better response than the 10-µg dose. Accordingly, a greater dose was used in further studies. Determining the optimal vaccine program, including the optimal timing and number of doses, requires further testing. Based on current results, at least two doses are required. From an economic perspective, further studies are also needed to determine whether a smaller vaccine dose or a combination of a smaller and larger dose would be sufficient.

Shuai et al showed that a spike-protein-based vaccine was effective in mink against the early virus strain(8). In our study, nAb titers were somewhat higher 5 weeks after vaccination. However, titers between studies are not fully comparable due to differences in protocols. IgG and nAb levels remained detectable over a 6-month follow up despite exhibiting a slow decline, as has been observed with other vaccines, including Moderna (mRNA-1272) and Pfizer (BNT162b2) in humans(26, 27). However, after infection, both IgG levels and nAb titers exceeded the upper detection limits in all vaccinated animals, whereas a smaller increase was observed in unvaccinated animals that were not been previously exposed to the virus. This indicates the presence of immunological memory in vaccinated mink despite the decrease in antibody levels and possible protection even several months after vaccination. A similar phenomenon has also been observed in humans; for example, a stronger antibody response against Omicron infection was detected in fully vaccinated individuals(28, 29).

Current SARS-CoV-2 vaccination strategies rely on intramuscular immunization (IM), which is insufficient to fully prevent transmission, possibly due to insufficient activation of mucosal immunity. Hence, intranasal (IN) vaccines have been suggested and several are undergoing clinical trials(30). For example, Hassan et al showed that a spike-protein-based chimpanzee adenovirus vector vaccine induced a stronger IgG response when given IN as compared with IM and, unlike IM vaccination, the IN route also induced an IgA response(31). Other suggested advantages of IN vaccines include a possible lower sufficient dose (leading to improved safety and lower costs), protection against other virus variants, easier administration, and lack of requirement for needles or a sterile environment(32). Here, the RBD vaccine was shown to induce an antibody response when given IN even though the response was smaller than the IM vaccine. However, this may be due to the lack of optimization regarding the correct dose and delivery mechanisms to avoid clearance by nasal barriers. Further tests should be conducted with different doses and adjuvants. In addition, comparing mucosal immunity (IgA) and performing a virus challenge in IN vaccinated mink is required.

A few 0-samples (3%, 6/191) taken before vaccination had an elevated signal (0.376-0.931) in ELISA. These were all tested with MN, in which no NAbs were detected. These samples were considered false positives. The reasons for false positives remain unclear, but one possibility is cross reaction with some host factor or mink coronavirus. This should be considered when using ELISA in SARS-CoV-2 diagnostics. Positive results, especially from clean farms and households, should always be confirmed with other methods to avoid reporting false positive results.

Our results indicate that other vaccinations may hinder the SARS-CoV-2 immune response if given simultaneously. Although no statistically significant effect was seen with the larger dose, this possibility cannot be excluded with this sample size. A statistically significant difference was observed with a smaller dose, which suggests that the Febrivac-3 plus should not be given simultaneously with SARS-CoV-2 vaccine. In addition, other vaccine combinations also require testing before considering simultaneous administration. In humans, this relationship has not been observed between SARS-CoV-2 vaccine and influenza vaccine(33). However, the possibility of this phenomenon with other vaccine combinations cannot be excluded. It should be noted that we only tested the SARS-CoV-2 response. To determine if the SARS-CoV-2 vaccine disturbs the immune response against the pathogens included in Febrivac 3-Plus, specific antibody tests should be performed.

Consistent with results from earlier studies(21, 34), virus was shed in saliva of vaccinated and unvaccinated mink from 1 dpi. However, comparison of Ct values indicates that the amount of viral RNA was smaller in vaccinated mink, which suggests a possible lower viral load in saliva, consistent with results from humans after vaccination with BNT162b2(35). Virus was cultured from saliva of unvaccinated mink for 6 days and for only 4 days from the saliva of vaccinated mink. This contrasts with the results of Shuat et al., who did not culture the virus from vaccinated mink(8). However, their sample materials were different from ours and sample size was smaller (three vaccinated animals). They also used a different virus variant isolated from a patient in the early phase of the pandemic, whereas our study was performed with an Alpha variant, which has been shown to spread more easily in humans(36). A notable difference between variants is that the virus was cultured from saliva for a longer time in Alpha-infected mink than in Omicron-infected mink. Based on the naturally infected recipients, virus was cultured from saliva even before saliva was PCR positive.

The clinical signs were like those observed by Adney et al.(34). Comparison of clinical signs suggests that vaccinated mink had less nasal irritation and possibly slightly less anorexia, diarrhea, malaise, and eye symptoms. However, it should be noted that evaluation of clinical signs is challenging and uncertain and many of the signs can also be caused by stress due to change of environment. Different caretakers may also evaluate the signs differently, which is why overall health of the animals was always evaluated by the same person. Despite the uncertainty of evaluating clinical signs, histology results strongly support the conclusions of milder disease in vaccinated mink. Surprisingly, mink infected with Omicron appeared to have more clinical signs than mink infected with Alpha. However, the experiment with Omicron was performed later, after more information about SARS-CoV-2 in mink was available, which is why we were better able to evaluate the clinical signs. Further studies are needed to confirm these differences between clinical signs.

In humans, men have been suggested to be more susceptible to severe COVID-19 than women(37). Women also have a more robust vaccine response than men(38). Accordingly, we also compared male and female mink. As many ELISA values were above the upper limit of detection, differences in IgG response could not be reliably assessed. However, smaller amounts of nAbs were detected in male mink in all vaccine groups. Although the results were not statistically significant, the sample size of only 14-15 animals per group should be noted. In the virus challenge with the Alpha variant, male mink appeared to have more clinical signs than female mink, although no difference in Ct values from saliva were observed. The same difference was not observed with Omicron, although the smaller sample size should be noted. Before infection, mink exhibited some signs of anorexia, diarrhea, and very mild nasal and eye irritation. However, the condition of these mink clearly deteriorated after infection and more signs appeared, indicating that Omicron caused a more severe clinical disease in both male and female mink.

The neurotropism of SARS-CoV-2 has been reported in human as well as in experimental animal models(39–41). Despite these reports, the neuropathogenesis of SARS-CoV-2 is not well understood, especially in the post-infectious context. Repercussion of vaccination on viral dissemination to the central nervous system (CNS) is even more poorly known. Here, we showed the presence of SARS-CoV-2 in CNS of unvaccinated mink. Since viral infection of CNS has been suggested as one of the potential reasons of neurological symptoms associated with long COVID(42), mink could also serve as an animal model for long COVID. For this, further studies with longer follow-up period and bigger emphasis on behavioral symptoms should be performed. Additionally, we demonstrated that vaccination reduced virus entrance into the brain. This is in line with reports in humans as several studies have indicated that vaccination may decrease the risk for long COVID(4).

SARS-CoV-2 has a broad host spectrum and has potential to infect new host species. This increases the risk of formation of virus reservoirs and generation of new, possibly more dangerous variants. In addition to humans, symptomatic disease has also been described in cats, tigers, lions, and mink(18, 43–45). Due to its zoonotic and reverse zoonotic potential aiding in the development of new variants, it is also important to study and control the virus in other host populations. RBD mFc-based vaccines may have potential for use not only in humans but also in domestic animals, due to indications of milder disease and lower viral load. Even though the vaccine tested here most likely does not fully prevent further infections (like in current human vaccines), it is likely to decrease the risk of spread and improve animal welfare. The viral load used in this experiment is likely greater than the actual viral exposure in real-life situations, meaning that the protective effect in real life may be greater than that observed here. A study including experimentally infected animals and uninfected, vaccinated recipients should be conducted to determine if the vaccine prevents infection of vaccinated animals.

## Materials and methods

### Protein and vaccine production

RBD with or without mouse Fc was produced as previously described(46). The vaccines were prepared by diluting the purified proteins to set concentrations and then adding the adjuvant to one third of the final injection volume (1 ml) and vortexing for 30 s. This suspension was kept at 4°C until time of injection.

### Ethical permit, vaccination and sampling

All the experimental procedures were approved by the Animal Experimental Board of Finland (ESAVI/33259). In the first experiment, 6-month-old brown (mahogany) mink (30-31/group, Fig. 6) were vaccinated subcutaneously with 40 µg of RBD mFc (RBD mFc big), 20 µg of RBD mFc (RBD mFc small), 10 µg of RBD (RBD small), or adjuvant only (adjuvant) or intranasally with 40 µg with RBD mFc (intranasal). A second dose was given 2 weeks after the first dose. Vaccinations were done either concurrent with sampling or to unsedated animals in the sheds. Subcutaneous vaccinations were administered in the inguinal skin fold. The final volume of vaccinations was 1 ml. Mink from RBD mFc small and RBD small groups were euthanized after 11 weeks, whereas the remainder were kept alive after the experiment. Five mink died during the follow up in the RBD mFc big, 3 died in the RBD mFc small, 1 died in the RBD small, 4 died in the intranasal, and 2 died in the adjuvant groups. This was considered a normal loss of animals and upon independent necropsy by the Finnish food authority pathologists the causes were deemed unrelated to the vaccine.

**Fig. 6.**
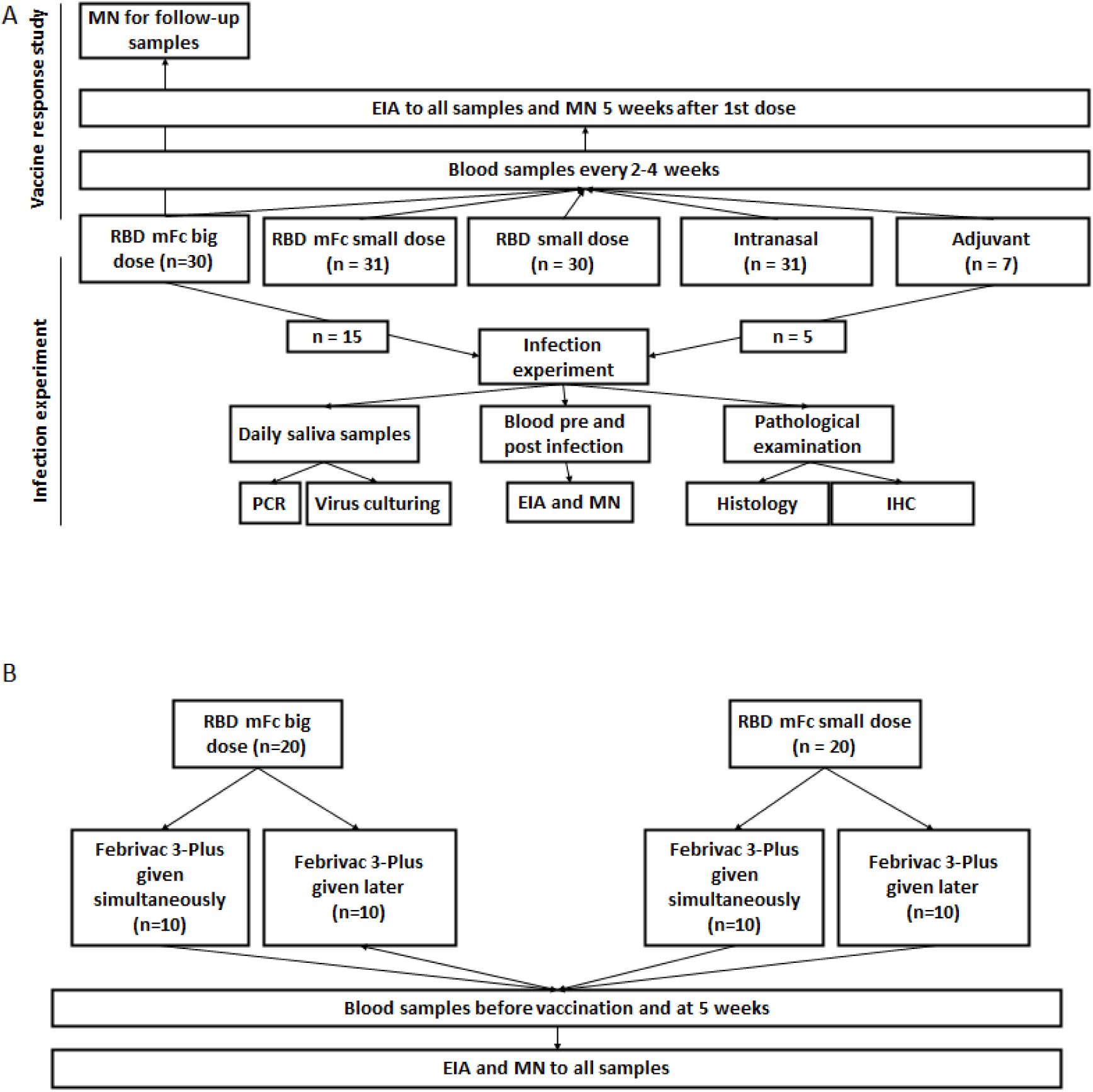
Flowchart of experiments 1 and 2.s

Blood samples were taken before vaccination and every 2-4 weeks for up to 27 weeks. Before sampling, the animals were anesthetized with Domosedan (detomidine hydrochloride 10 mg/ml) and Ketaminol (ketamine hydrochloride 50 mg/ml). The dosage was 0.1 mg detomidine hydrochloride/kg and 5 mg ketamine hydrochloride/kg intramuscularly (IM). A blood sample of 2 ml was taken from the jugular vein with open-needle technique. After sampling, anaesthesia was reversed with Antisedan (atipamezole hydrochloride 5 mg/ml) 200 µg/kg IM. Euthanasia was performed with a triple dose of detomidine hydrochloride and ketamine hydrochloride and 0.35 ml/kg of intracardial Euthasol (pentobarbital 400 mg/ml)

In the second experiment, 20 mink were vaccinated with RBD mFc big and 20 with RBD mFc small as described previously. Blood samples were taken before vaccination and 5 weeks after the first dose. Ten mink from both groups received Febrivac 3-Plus vaccine at the same time as the first dose and 10 mink received Febrivac 3-Plus 5 weeks after the first dose (3 weeks after the second dose).

### Cell line and virus strains

Vero E6 cells (VE6) and their TMRPRSS2-expressing clone (VE6T)(47) were grown in minimal essential Eagle’s medium (MEM, Sigma-Aldrich) with fetal bovine serum (FBS, ThermoFisher, 10% for cell maintenance and 2% for infection), 2 mM L-glutamine, 100 IU/mL penicillin, 100 g/mL streptomycin, and 0.205 μg/ml of amphotericin B (Fungizone, Thermo Scientific). Alpha variant (B.1.1.7)(48) was used in the experimental infection. The following virus strains were used in MN: SARS-CoV-2/Finland/1/2020 (FIN-1, VE6-adapted strain from first case in Finland), and wild-type strains C1P1, Alpha (VoC1, B.1.1.7), Beta (VoC2, B.1.351), Delta (26), and Omicron BA.1 (OM393712)(48–50).

### Antibody tests

SARS-CoV-2 ELISA was performed on all blood samples. First, 96-well plates were coated at 4°C overnight with RBD (described previously) diluted in PBS (50 µl/well, 12 µg/ml) and washed three times with 250 µl PBS with 0.1% Tween 20 (TPBS). Wells were then blocked with 200 µl of 3% skim milk + TPBS for 1 hr at RT and washed as before. Serum (100 µl/well) diluted 1:200 in 1% skim milk + TPBS was added, followed by 2 h incubation at RT and washing as before. Plates were incubated for 1 h at RT with 50 µl of goat anti-ferret IgG (H + L) secondary antibody (Abcam UK) diluted 1:12 000 in 1% skim milk + TPBS and washed as earlier. Results were visualized by incubating with TMB (100 µl/well) for 15 min and followed by addition of 100 µl 0.5 M H_2_SO_4_. A450 was measured with a VICTOR^3TM^multimeter.

Microneutralization tests were performed according to the protocol by Haveri et al.(50) using a two-fold dilution series from 1:20 onwards. Samples from vaccinated mink were titrated until the correct titer was acquired and samples from experimentally infected mink until 1:2560. VE6 cells were used for FIN-1 and VE6T cells for wild-type strains. From the first experiment, blood samples of mink vaccinated with RBD mFc big from nine timepoints, samples from other groups from the 5-week timepoint, pre-infection blood samples, and terminal samples were tested with FIN-1. Five-week timepoint samples of RBD mFc big were also tested with wild-type C1P1, Alpha, Beta, and Delta variants. From the second experiment, all blood samples were tested with FIN-1. The RBD mFc big group that received Febrivac 3-Plus 5 weeks into the experiment was also tested with Omicron. Results from all antibody tests from the first experiment are shown in Table S1 and from the second experiment in Table S4.

### Infection experiment with alpha variant

Fifteen mink vaccinated with RBD mFc big were selected for virus challenge. Five unvaccinated mink from the group that received only adjuvant were also selected as unvaccinated controls. Mink were first transported from the research farm to Helsinki University test animal center for an adaptation period of 2-4 weeks wherein they were able to accustomate to the custom made cages with nesting boxes and non-invasive saliva sampling methods. This period was also used to educate personnel working with the animals on signs of stress or disease, handling, feeding, and sampling. Mink were transported into a BSL-3 laboratory and acclimatized for 2 days. For infection, animals were anesthetized as described earlier and inoculated with 2.5 ml of 8×10^5^ IU/ml virus intranasally. Blood samples (Copan) were collected before infection. Health of the mink was assessed daily by a professional animal caretaker (always the same person, blinded to which animals were vaccinated) who assessed the animals for sneezing or runny nose, diarrhea, anorexia, cough, eye symptoms, lethargy, and other clinical signs. Clinical signs were scored 1-3 (1 = mild, 2 = moderate, 3 = severe) and numbers of each symptom in each animal were recorded. Saliva samples were collected every day after infection with foam swabs (Virocult, MWE) into 200 µl of PBS and 200 µl of cell culture media (2% FBS). After 7 dpi, animals were anesthetized intramuscularly with detomidine (0.15 µg/kg) and ketamine (7.5 mg/kg) and euthanized in a CO_2_ chamber. Under terminal anesthesia, an intracardiac blood sample (2 ml) and a salivary sample were collected. After euthanasia, head, trachea, heart, lungs, liver, spleen, kidneys, intestinal tract (stomach, duodenum, jejunum, and colon), pancreas, bladder, adrenals, thyroid glands, skeletal muscle, mediastinal and mesenteric lymph nodes, and bone marrow were sampled for pathology or PCR. The Animal Experimental Board of Finland approved the experiment (ESAVI 33259).

### PCR and cell culturing of alpha-variant infected mink

All saliva samples were cultured in VE6T cells for 7-10 days and monitored for signs of CPE. If CPE was visible or there was excess fungal growth in the sample for accessing CPE, cell culture media was collected for PCR.

RNA was extracted from saliva and cell culture samples with a Viral RNA minikit (Qiagen) according to the manufacturer’s instructions. Extracted RNAs were tested with RdRp-targeting RT-qPCR(51). Cell culture samples were considered positive for virus growth if the Ct value from culture media was over 5 cycles lower than that of the original saliva sample, a possible positive if it was 1-5 cycles lower, and negative if it was similar or greater. All PCR reactions were performed with Stratagene Mx3005P instrument (Agilent Technologies).

### Histopathological examination

The head, trachea, heart, lungs, liver, spleen, kidneys, intestinal tract (stomach, duodenum, jejunum, and colon), pancreas, bladder, adrenals, thyroid glands, skeletal muscle, mediastinal and mesenteric lymph nodes, and opened femur (for sampling bone marrow) were fixed in 10% neutral-buffered formalin. After 48 h fixation, organs were cut and fixated for a further 24 h. Lung samples covered left and right cranial and caudal lobes and right medial lobe. A sagittal nasal cavity sample, including the vestibular, respiratory, and olfactory region of nasal conchae, was decalcified in 14.3% EDTA (pH 7.0). The fixed samples were processed routinely, embedded in paraffin, sectioned at 4-μm thickness, and stained with haematoxylin and eosin (HE). Histopathological scoring was based on lesion severity and the extent/proportion of the affected tissue. The lesions detected were scored as 0 = absent, 1 = mild, 2 = moderate, and 3 = severe. The total score of lung lesions was based on severity of bronchitis/bronchiolitis, interstitial pneumonia, alveolar damage, vascular lesions, and degree of pneumocyte type II hyperplasia. A total score of 11-15 was considered as severe, 6-10 moderate, and <5 as mild lesions. Histopathological findings in the other organs were also recorded.

### Immunohistochemistry

Paraffin-embedded nasal cavity and lung samples were immunohistochemically (IHC) stained for SARS-CoV-2 antigens employing custom-made rabbit polyclonal anti-SARS-CoV-2 spike S1 RBD and anti-SARS-CoV-2 nucleoprotein (rec. NP) antibodies (Abs). In addition, selected samples were subjected to IHC staining with the following Abs: MAC387 (dilution; cat no, AbdSerotec,) for macrophages, CD79a (dilution; cat no, AbdSerotec) for B lymphocytes, and CD3 (dilution; cat no, Dako) for T lymphocytes.

For the SARS-CoV-2 IHC, deparaffinized slides were incubated for 20 minutes at 99°C in 10 mM citrate buffer (pH 6) for antigen retrieval. Endogenous peroxidase was quenched with 3% hydrogen peroxide and nonspecific Ab binding blocked with 10% bovine serum albumin in PBS. The primary Abs, rec NP and RBD, were incubated for 1 h at room temperature diluted (rec NP, 1:3000; RBD, 1:1000) in an animal-free blocker and diluent solution (R.T.U. Animal-Free Blocker and Diluent; Vector Laboratories, Burlingame, Ca, USA). The secondary Ab, polymer-linked to HRP (BrightVision + Poly-HRP kit; ImmunoLogic, Duiven, Netherlands), was incubated for 30 minutes at room temperature and visualized with Bright DAB Substrate kit (ImmunoLogic, Duiven, Netherlands). Subsequently, sections were counterstained with Harris haematoxylin, dehydrated in ethanol and xylene, and cover slipped with Pertex® mounting medium (HistoLab).) The primary Ab was replaced with Rabbit IgG Isotype control (1:1000) for negative control slides.

### *In Situ* Hybridization

*In situ* hybridization (ISH) was performed using RNAscope® technology (Advanced Cell Diagnostics, Newark, CA) using SARS-CoV-2 RNA S-gene probe encoding for the spike protein (RNAscope™ Probe-V-nCoV2019-S-C3, Cat No. 848561-C3) and RNAscope 2.5 HD Reagent Kit-Red (cat. 322350). The protocol used strictly followed the manufacturer’s protocol without any modifications. Internal ISH method positive control included formalin-fixed paraffin-embedded (FFPE) sample from mouse liver and the probe mouse Mm-Ppib (catalog number 313911, ACDbio). Positive control for SARS-CoV-2 included an FFPE sample of mink lung with SARS-CoV-2 positive immunohistochemistry results. Non-specific tissue hybridization was tested by using the probe mouse Mm-Ppib (catalog number 313911, ACDbio) on a FFPE brain sample from one mink.

### Infection experiment with Omicron variant

We previously experimentally infected 5 male American mink with the Omicron variant. The experiment was repeated with 5 female mink with the same experimental setup as described previously(21). Briefly, 3 mink were infected and 2 were left as uninfected recipients. Experimentally infected mink were followed for 7 days and recipients for 10 days. Saliva samples were taken and tested with PCR (Luna SARS-CoV-2 RT-qPCR Multiplex Assay Kit (NEB)) and cell culturing daily. Histopathological studies were performed after the experiment.

### Statistical analysis

Analysis was performed with IBM SPSS Statistics 28 or RStudio utilizing dplyr, psych, ggplot2, and ggpubr packages(52–57). Microneutralization values below 20 were set to 10 for calculations and PCR samples with no Ct were set to 40. The data were tested for normality using Shapiro-Wilk test. Parametric tests (independent samples T test [with Levene’s test and two-sided p-value]) were used for normally distributed data and non-parametric tests (independent samples Mann-Whitney *U* test, independent samples Kruskal-Wallis test, or related samples Friedman’s two-way analysis of variants by ranks test) for non-normally distributed data. Statistical significance level was set to 0.05. MN results were visualized in log_2_-scale and using dodge function. Jitter function was used to visualize ELISA results.

## Acknowledgements

We wish to thank Kati Kuipers, Anne Kujanpää and Laura Vähälä from Finnish Centre for Laboratory Animal Pathology for building up and performing immunohistochemistry and in situ hybridisation. Warm thanks also to animal care takers Jari and Mari Elemo and Oleksandr Cherniavskyi for their help tending the animals and Esa Pohjolainen and Olga Kivelä for technical assistance.

## Author contributions

KA, JV, RK, LK, HN, AS, OR, JK, JP, and TS designed the study; OR, RN, and AP developed and produced the antigen used for the preliminary version of the vaccine; JK, SS, TH, MI, MIO and EO implemented the vaccine trial; TN and KA did the serological tests; JV, KA, RK, EK, RM, VV, and LK performed the virus challenge and sampling; VV and TN performed the PCR tests; KK, HN, and JL participated in pathological analysis; JV performed the statistical analysis; JV, KK, and HN handled the visualization; KA, RK, JP, and TS supervised the study; JV, HN, KK, and KA wrote the original draft; all authors helped revise the manuscript.

## Data availability statement

All the data is available in the manuscript or its online supplementary material.

