## Supplemental figures and methods for "American mink as an animal model to study SARS-CoV-2 and vaccine response"

### Supplementary data

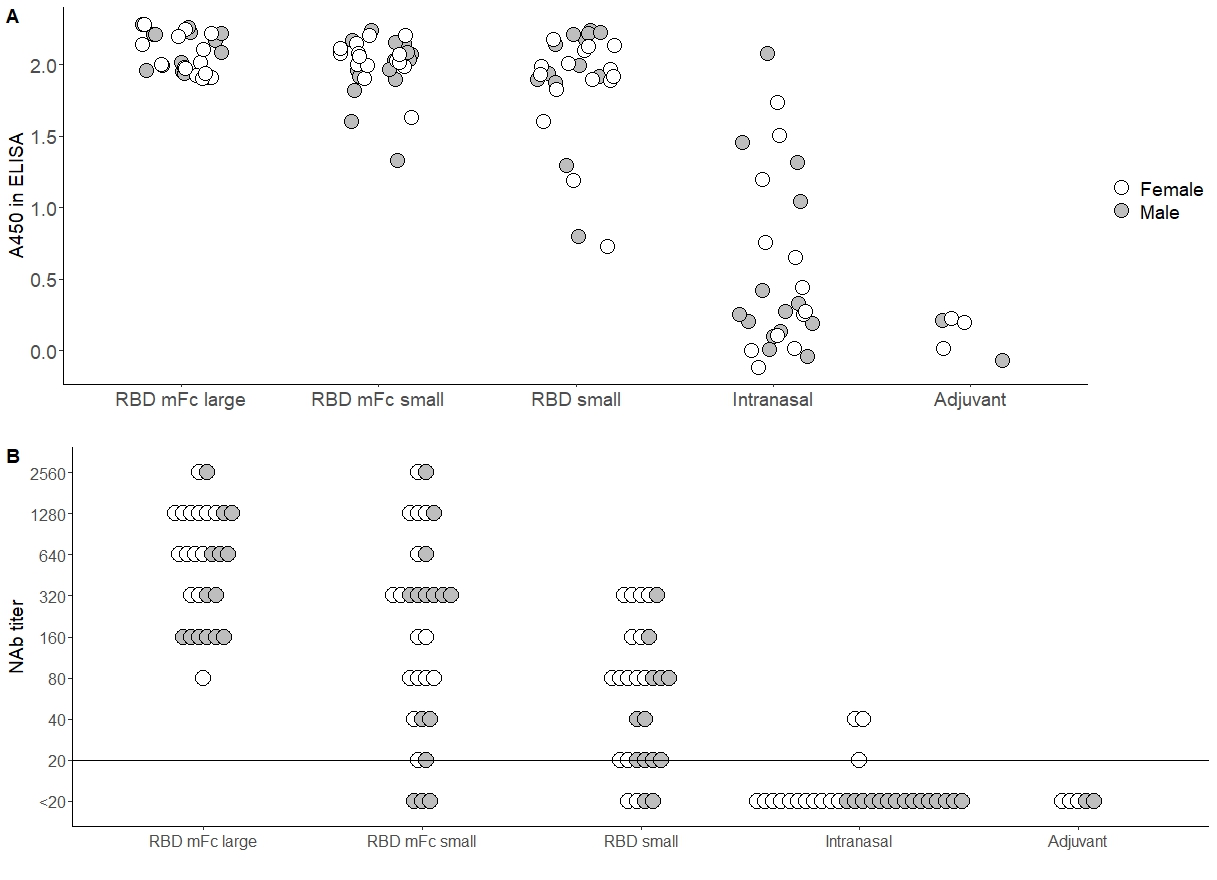

Fig. S1: IgG levels (a) and NAb titers (b) five weeks after the first and three weeks after the second dose in SARS-CoV-2 vaccinated mink separated by gender. Highest detection limit is 2.0-2.1 in ELISA. NAb-titers are expressed in log_2_ scale and limit of detection marked with horizontal line.

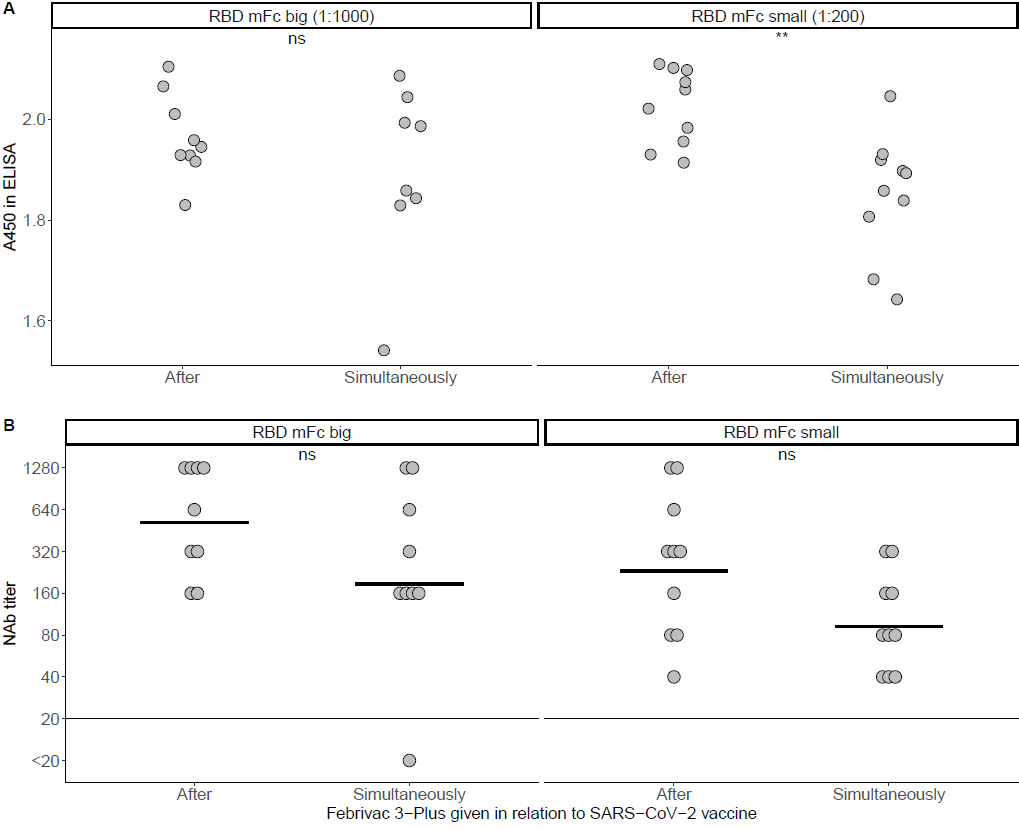

Fig. S2: IgG levels (a) and NAb titers (b) in mink that received Febrivac 3-Plus at the same time or five weeks after the first dose of SARS-CoV-2 vaccination. NAb titers are expressed in log_2_ scale and ELISA upper detection limit is 2.0-2.1. Limit of detection in MN is indicated with a horizontal line and statistical significances (**: < 0.01, ns: not significant) are indicated above the pictures.

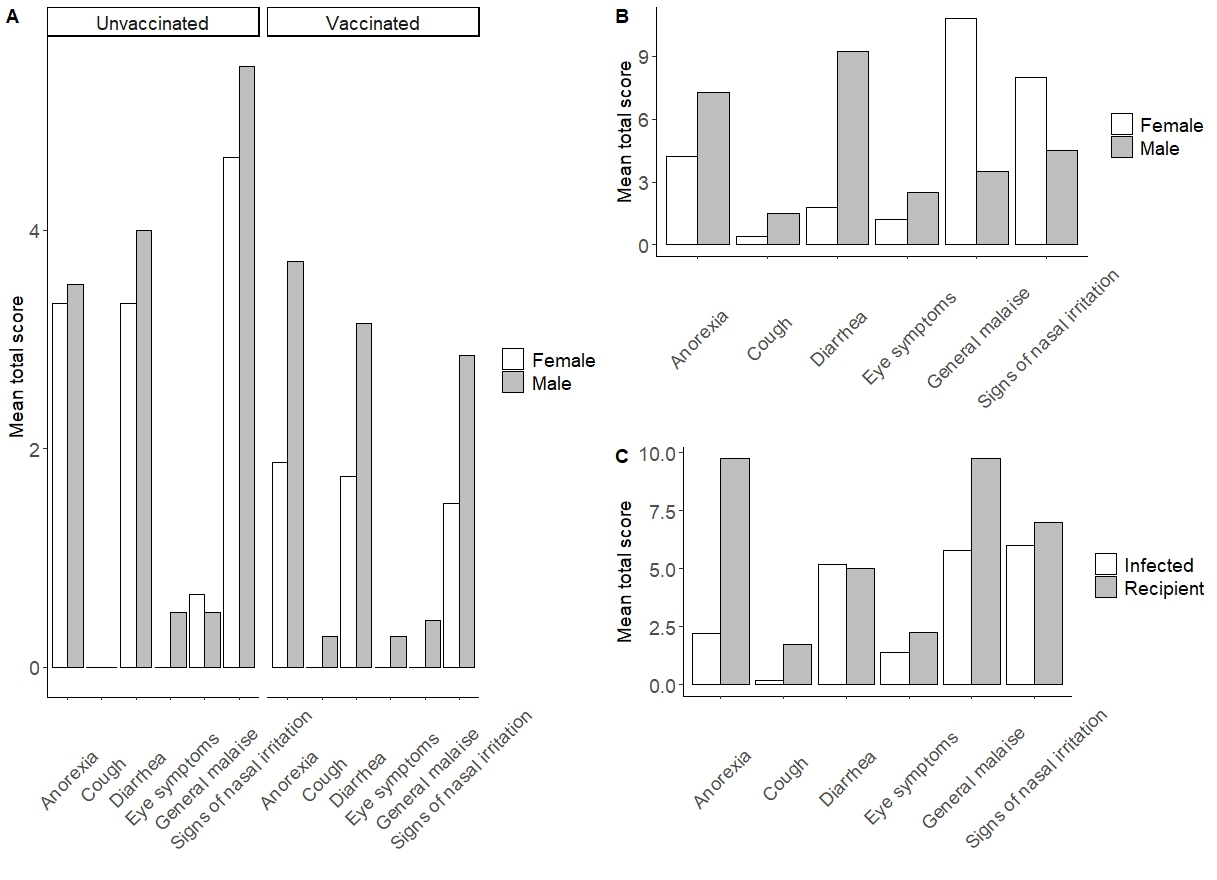

Fig. S3. Clinical signs in mink experimentally infected with alpha (a) variant and omicron (b and c) variant. Clinical signs of each class were scored 0-3 (0 = none, 3 = severe) daily and sum with all the days combined was calculated (=total score). Anorexia and gastrointestinal symptoms may also result from a stress of changing the environment.

**ELISA-test validation and analysis for SARS-CoV2 in mink**

The ELISA-test used in this study was modified from Amanat et al. 2020 assay for detecting SARS CoV-2 seroconversion in humans. The test was validated by testing 24 SARS CoV-2 negative mink serum samples and 4 pools of negative mink serum with 8-10 (2 x 8 and 2 x 10) sera in each pool so altogether 62 negative mink sera. The sera tested were both AMDV positive (4 single sera and 1 pool) and AMDV negative (20 single sera and 3 pools). The serum samples were collected during 2008 to 2017 for Aleutian mink disease diagnostics and had been preserved at -80 °C. The cut off was calculated as the mean plus 2x standard deviation and based on these results was 0.3. The analysis was further tested by including 10 SARS CoV2 positive mink blood samples obtained from Dr Harold van der Heijden and DMV Robert Jan Molenaar, taken during the primary epidemic in The Netherlands. We found quite a large percentage (10-20%) of false positive ELISA results from our initial tests with full spike protein and N protein as well as the plates used by the original authors. The protocol was modified to use the SARS-CoV-2 RBD (Wuhan) protein and the plate used in the analysis was changed to MaxiSorp (Thermo-Fiscer Scientific, USA), and the sample dilution was raised to 1:200 from the original 1:100. The number of false positive results was thus lowered to approximately 1 % of all tested samples. If ELISA tests were to be used for SARS-CoV-2 diagnostics, a second sample would be required to show diagnostic rise in antibody titers. Our initial tests on the experimental animals included in this study showed that when samples were taken two weeks apart the false positive animals tested negative in the later sample. No connection between sample quality such as hemolysis or lipemia could be shown to have an effect on the quality of the analysis. Nor did the AMDV status of the animals seem to affect the test.
